# DeepKinZero: Zero-Shot Learning for Predicting Kinase-Phosphosite Associations Involving Understudied Kinases

**DOI:** 10.1101/670638

**Authors:** Iman Deznabi, Busra Arabaci, Mehmet Koyutürk, Oznur Tastan

## Abstract

Protein phosphorylation is a key regulator of protein function in signal transduction pathways. Kinases are the enzymes that catalyze the phosphorylation of other proteins in a target specific manner. The dysregulation of phosphorylation is associated with many diseases including cancer. Although the advances in phosphoproteomics enable the identification of phosphosites at the proteome level, most of the phosphoproteome is still in the dark: more than 95% of the reported human phosphosites have no known kinases. Determining which kinase is responsible for phosphorylating a site remains an experimental challenge. Existing computational methods require several examples of known targets of a kinase to make accurate kinase specific predictions, yet for a large body of kinases, only a few or no target sites are reported. We present DeepKinZero, the first zero-shot learning approach to predict the kinase acting on a phosphosite for kinases with no known phosphosite information. DeepKinZero transfers knowledge from kinases with many known target phosphosites to those kinases with no known sites through a zero-shot learning model. The kinase specific positional amino acid preferences are learned using a bidirectional recurrent neural network. We show that DeepKinZero achieves significant improvement in accuracy for kinases with no known phosphosites in comparison to the baseline model and other methods available. By expanding our knowledge on understudied kinases, DeepKinZero can help to chart the phosphoproteome atlas.

## 1 Introduction

Protein kinases are a large family of enzymes that catalyze the phosphorylation of other proteins [1]. Phosphorylation involves the transfer of a phosphoryl group to the side chain of an amino acid residue in the substrate. The amino acid residue that receives the phosphoryl group is called the phosphorylation site, or briefly a *phosphosite*. The phosphosite is usually one of the three amino acids, serine, threonine, and tyrosine although other amino acids, such as histidine, are being reported to act as phosphosites [2]. Phosphorylation events can lead to the activation or deactivation of proteins, modify the targets’ interactions with other proteins, direct them to sub-cellular localization or target them for destruction [3]. Since they are the key regulator of protein function in a broad range of cellular activities, aberrant kinase function is implicated in many diseases [4], particularly in cancer [5, 6]. Several pathogenic human mutations also lie on known phosphorylation sites [7]. Kinases, therefore, are also major drug targets [8, 9]. To this end, understanding the associations between kinase and phosphorylation sites holds the key to understanding the signaling mechanisms in the healthy and diseased cells.

Advances in mass spectrometry-based phosphoproteomics has enabled identification of phosphosites at the proteome level [10–12]. Many computational models have also been developed to predict phosphosites in a given input protein sequence (recently [13–20] and earlier methods reviewed in [21]). Once a phosphosite is identified, either experimentally or computationally, determining the kinase that is responsible for catalyzing the phosphorylation of this site becomes the key question. With 518 identified kinases in the human genome [22] and the transient nature of kinase-substrate interactions, it is still an experimental challenge to determine the kinase that targets a given site. As underlined by a recent review [7], most of the phosphoproteome is uncharted: more than 95% of reported human phosphosites have no known kinase or associated biological function.

To identify a phosphorylation site on a protein sequence, several computational methods have been proposed earlier [13, 15, 17–20, 24–32]. Since these methods can also provide kinase specific predictions, they can be used to predict associated kinases of a known phosphosite. A majority of these methods utilize consensus sequence motifs or position specific scoring matrices to estimate the position preferences of each kinase. This approach requires a reasonable number of previously known targets to be able to estimate the positional preferences of a kinase accurately. Other tools employ supervised machine learning models that use a collection of established kinase-phosphosite associations. They model the relationship between the properties of kinases and the properties of their target phosphosites in a supervised classification setting. The application of such tools is limited to kinases for which a substantial number of target phosphosites are available for training. For example, MusiteDeep [20], uses deep learning to predict binding sites for kinases, and it exclusively focuses on kinase families with at least 100 experimentally verified phosphosites. Recently, the use of phosphorylation data to predict kinases has been proposed, but these methods also require knowledge of target sites for a kinase to make predictions for that kinase [33]. Some of the recently developed tools and the number of kinases and/or kinase families they predict are shown in Table 1 along with the number of sites required for a kinase to be included.

**Table 1:** Kinase coverage of state-of-the-art sequence-based methods for predicting kinase-substrate associations. For each method, the middle column reports the number of kinases and kinase families for which the method can predict target phosphosites. The right column reports the criteria employed by the method for being able to make predictions for a kinase or family.

| Method | Number of kinases or kinase families | Criteria for inclusion |
| --- | --- | --- |
| MusiteDeep [20] | 5 families | Families with $> 100$ sites |
| PhosphoPredict [18] | 8 families | Families with $\geq 50$ sites |
| Li et al. [23] | 8 families | Families with $\geq 50$ sites |
| PhosphoPICK [15] | 59 human kinases | Kinases with $> 10$ sites |
| PKIS [24] | 56 human kinases | Kinases with $> 10$ sites |
| KSRPred [17] | 103 human kinases | Kinases with $\geq 15$ sites |
| KinomeExplorer [13] | 222 kinases covered but accuracy assessed for 14 kinases | Kinases with $\geq 20$ sites |

A particular problem that has been overlooked in the literature is the prediction of target phosphosites for kinases with few or no known phosphosites. Despite the central role of kinases in cellular signaling cascades and their importance as potential drug targets, a large fraction of the kinome is understudied [7, 9, 34]. PhosphositePlus, a database of experimentally validated phosphosites, provides phosphosite annotations for only 364 human kinases. For nearly 200 of 364 annotated kinases, there are at most 10 experimentally validated target sites (Figure 1).

**Figure 1:**
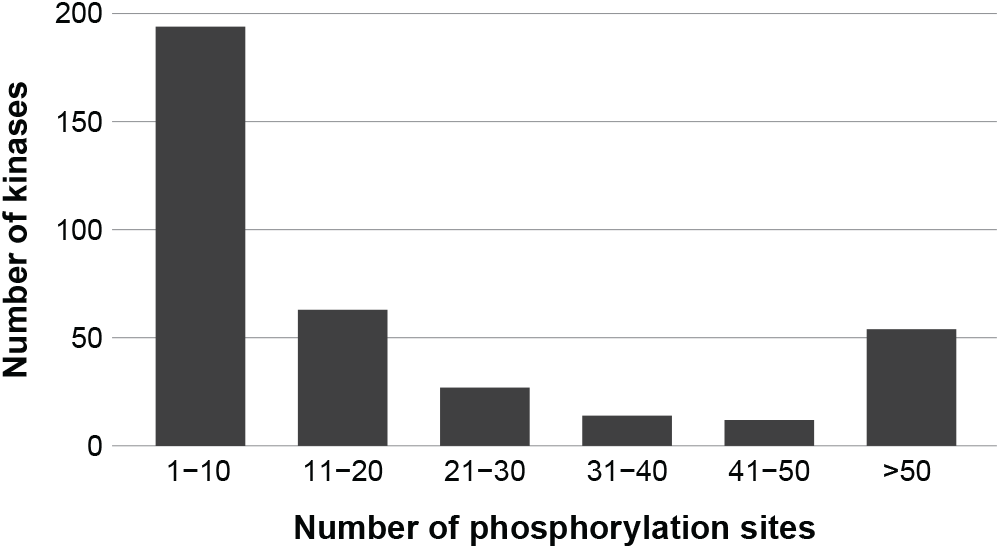
The distribution of the number of experimentally validated target phosphosites for kinases in the human kinome. The histogram is based on data obtained from PhosphositePlus database experimentally validated phosphosite-kinase interactions.

In this study, we introduce DeepKinZero, a zero-shot learning approach to predict kinase-substrate associations for kinases with no known target sites. Zero-shot learning is a machine learning approach that has received significant attention, particularly in the field of computer vision. It handles recognition tasks for classes for which no training examples are available [35–39]. The key to making predictions for classes with no training data (referred to as *unseen* or *zero-shot* classes) is to have side information which can be used to relate the classes. Based on these relations it becomes possible to transfer the knowledge obtained from classes that have positive training samples (referred to as *seen* class) [39] to the previously unseen classes.

As exemplified by Yu et al. [40], it is difficult for an image classification system to recognize an okapi when there are no images of okapi in the training set. Yet, if the visual descriptions such as – zebra-stripes, four legs, brown torso, a deer-like face – can be learned from the seen classes (zebra, deer, horse, etc.) and if the system has side information indicating that okapis have these attributes, it becomes possible for the algorithm to recognize an okapi even without any prior exposure to an okapi visual. This is accomplished by detecting these visual descriptors and relating these descriptors to the side information on okapis. Similarly, even if we do not know any phosphosites that are associated with an understudied kinase (unseen class) in training, the zero-shot learning framework enables us to recognize a target site of this kinase by transferring knowledge from well-studied kinases to the rare kinases. This can be achieved by establishing a relationship between the kinases using relevant auxiliary information, such as functional, sequence and structural characteristics of kinases. It is important to note that, in the application of zero-shot learning to the prediction of kinase-substrate associations, phosphosites are represented as “instances” and kinases are represented as “classes” (i.e., kinase predictions are made for a given phosphosite). This is indeed the set-up that is relevant in many practical applications since the researchers who experimentally identify a phosphorylation site are interested in identifying kinases targeting that phosphosite.

Given a predicted or experimentally identified phosphosite, DeepKinZero predicts the most likely zero-shot kinase that can phosphorylate this particular site by using the local protein sequence centered at this site. DeepKinZero learns the phosphosite sequence features via a bi-directional recurrent neural network. Therein the kinases are represented based on functional and sequence information. Through learning a compatibility function that establishes relationships between the representations of the phosphosite sequences and the kinases, DeepKinZero transfers knowledge from kinases with many known phosphosites to those kinases with no known sites. We also consider alternate representations of the phosphosite sequence and the kinase embeddings, and asses their effectiveness. For kinases with no known target sites (i.e, kinases for which it is not possible to make predictions using other supervised methods), DeepKinZero provides predictions with 30-fold increase in accuracy as compared to random guess.

DeepKinZero offers a scalable and flexible approach annotating sites with kinases with no prior information on target sites. It is implemented in Python with the help of Tensorflow library [41] and is provided as an open source tool at https://github.com/Tastanlab/DeepKinZero.

## 2 Methods

### 2.1 Problem Formulation

The residues flanking the central phosphosite is critical for kinase specificity [42]. Thus, the local sequence surrounding the phosphorylation site has been a common input in the computational prediction of kinase-phosphosite associations. In this study, we use sequences of 15 residues (i.e., 7 residues flanking on each side of the phosphosite in addition to the phosphosite) as input and we denote these as the phosphosite sequences. Lengths of 15 or shorter have been shown to be useful in previous approaches [21, 43, 44]. Let 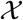 represent the space of phosphosite sequences and 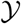 represent the set of all identified kinases in human. The problem of kinase-phosphosite association prediction is defined as follows: given a phosphosite sequence 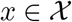, identify which kinase 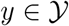 is most likely to catalyze the phosphorylation of this site. The problem is formalized as a multi-class classification problem with many classes, where each input phosphosite sequence is associated with a single kinase. This one-to-one mapping, in reality, does not always hold; a phosphosite occasionally can indeed be phosphorylated by more than one kinase. However, these cases occur rarely and in this study, whenever the predicted kinase is among the kinases known to phosphorylate a given phosphosite, we accept it as a true positive.

Some kinases are well-studied and many target sites have been identified for these kinases. On the other hand, many kinases lack formerly identified target sites. We refer to the kinases with known target sites in the training data as *common* kinases, these kinases constitute the training classes. We denote this set of kinases as 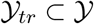. We call the kinases with few phosphosite annotation as *rare* kinases and denote the set of rare kinases as 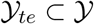. 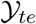 constitutes the zero-shot test classes. By definition, the sets of common and rare kinases are disjoint, i.e., *Y*_*tr*_ ∩ *Y*_*te*_ = ∅

The training data contains only pairs for common kinases, *D*_*tr*_ = {(*x*_*i*_, *y*_*i*_), *i* = {1, …, *N*_*tr*_}}, where 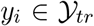. Since there are no positively labeled data for the rare kinases, 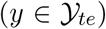, during the training phase, it is not possible to use traditional supervised methods to build a model for mapping sites to such rare kinases. However, it is known that some kinases are related to each other functionally, evolutionarily, or structurally [22]. Thus, using zero-shot learning, the known relationships between kinases can be exploited to learn a predictive model for rare kinases. In the next section, we elaborate on this approach.

### 2.2 The Zero-Shot Learning Model

Following the work by Akata et al. [39], we assume that a vector space representation, called class embedding or kinase embedding, can be constructed for each kinase. Therefore, an *m*-dimensional “kinase embedding” vector *ϕ*(*y*) ∈ ℝ^*m*^ can be computed for each kinase *y* ∈ *Y*. We expect similar classes to be close to each other with respect to the Euclidean metric in this embedded space. Similarly, for each phosphosite 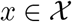, we compute the phosphosite embedding vector, *θ*(*x*) ∈ ℝ^*d*^, that represents the phosphosite sequence in a *d*-dimensional space. We discuss the computation of phosphosite and kinase embeddings in Sections 2.2.1 and 2.2.2 in greater detail.

#### The DeepKinZero Model

To accomplish transfer learning between the common and rare kinases, we learn the association between the phosphosite and the kinase embeddings. This idea is illustrated in Figure 2. Following the work in structured output prediction [45] and prior work in zero-shot learning [38, 39, 46–50], we use a compatibility function 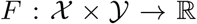 to model the mapping between the input and output embeddings. In this model, *F* takes a phosphosite - kinase pair (*x*_*i*_, *y*_*j*_) as input and returns a scalar value which is proportional to the confidence of associating the site, *x*_*i*_, with kinase *y*_*i*_. In this model, the probability that a given site is a target of a given kinase is calculated logistically from the bi-linear compatibility function *F*:

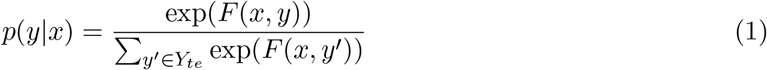

**Figure 2:**
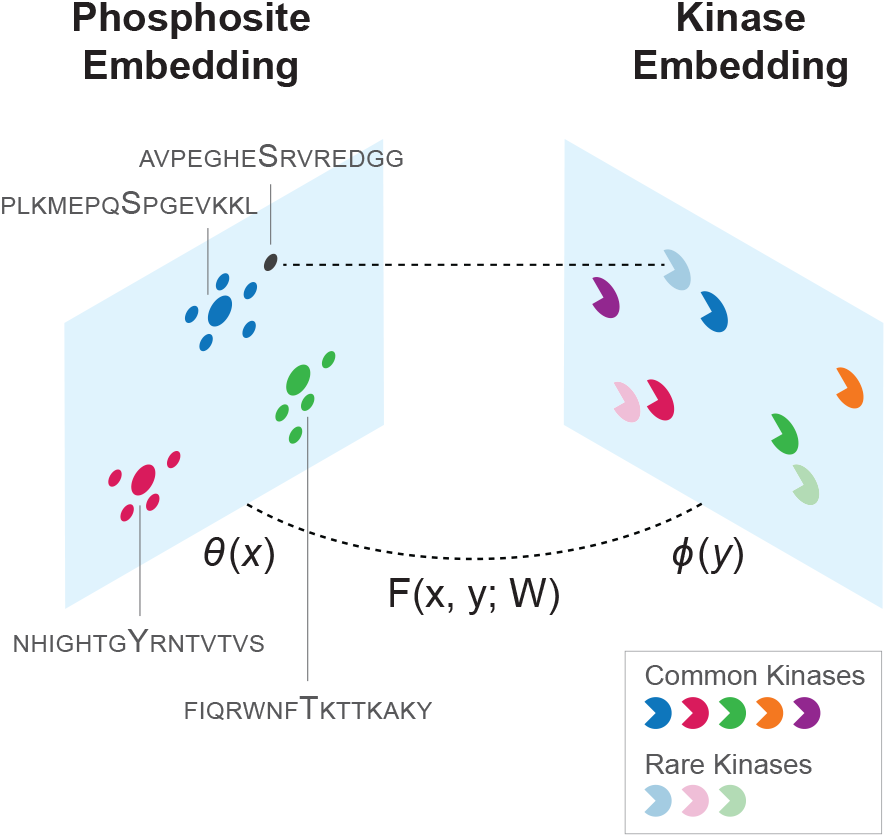
Overview of the application of zero-shot learning to the prediction of kinase-phosphosite associations. The phosphosites and the kinases are embedded into multi-dimensional vector spaces using the information on sites and kinases, respectively. The training data comprise common kinases and their sites. The parameters *W* of the function *F* (*x, y*; *W*) are estimated from the training data such that the compatibility between phosphosite embedding *θ*(*x*) and kinase embeddings *ϕ*(*y*) is maximized. For a new phosphosite at test time (shown as the black dot), the rare kinase that maximizes *F* for the input site’s embedding is picked by using *F* and the learned parameters 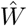

As in [50], we use the following bi-linear compatibility function for input *x* and *y*:

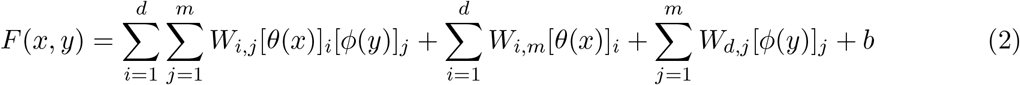

which can be written in matrix notation as:

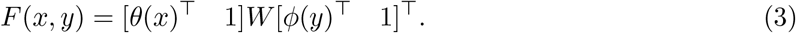

Here, [*θ*(*x*)]_*i*_ and [*ϕ*(*y*)]_*j*_ respectively denote the *i*-th and the *j*-th entries of the phosphosite and kinase embedding vectors, respectively. *W* denotes the (*d* + 1) × (*m* + 1) compatibility matrix, where *W*_*i,j*_ for 1 ≤ *i* ≤ *d* and 1 ≤ *j* ≤ *m* specifies the contribution of the correspondence between the *i*-th dimension in the phosphosite embedding space and the *j*-th dimension in the kinase embedding space to the compatibility of the phosphosite and kinase pair. *W*_*d*+1,*i*_ and *W*_*j,m*+1_ weights evaluate the information provided by the phosphosite and kinase embeddings individually. *W*_*i,m*+1_ for 1 ≤ *i* ≤ *d* specifies the weight of the *i*-th dimension in the phosphosite embedding space, *W*_*d*+1,*j*_ for 1 ≤ *j* ≤ *m* specifies the weight of the *j*-th dimension in the kinase embedding space. Finally, *W*_*d*+1,*m*+1_ = *b* denotes the bias term of the model.

We represent the 15-residue phosphosite sequences centering on each phosphosite with multi-dimensional vectors in Euclidean space, such that the embeddings of similar sequences are close to each other in this space. To learn phosphosite embeddings, we use Bi-directional Recurrent Neural Network (BRNN) [51] model with an attention mechanism over the training data. Recurrent Neural Networks (RNNs) constitute a class of neural networks that exhibit state-of-the-art performances for modeling sequential data [52]. At each time step, which corresponds to the current position in the sequence, RNN accepts an input sequence vector. The hidden state of the RNN is then updated via non-linear activation functions to predict the target class, which, in our case, is the associated kinase. BRNN contains 512 LSTM cells [53] on each direction. This number of cells is chosen to ensure the best compromise between memory requirements and accuracy performance on validation and training data.

We also employ a dot attention mechanism [54] over the output of the BRNN model to enable the model to focus on the more important positions of the input sequence. For this, we multiply the output vectors of BRNN with the attention vector *A*, which is *D* × 1. *D* is the size of the BRNN output embeddings, which is 1024 since we have 512 nodes on each side. Let *H* = [*h*_1_, *h*_2_, …, *h*_*T*_] denote the whole output of the BRNN. To calculate the attention value for each position, we multiply the attention vector with the output vector for the position *i*, denoted by *h*_*i*_. We apply softmax to the output of this multiplication to normalize them within the range 0-1. *α*_*i*_ = softmax(*h*_*i*_*A*). *α*_*i*_ is the attention weight for position *i*. Finally, the phosphosite embedding vector *ϕ*(*x*), is the weighted average of the positions by the attention weights: 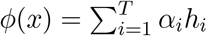

#### Training DeepKinZero

Given training data *D*_*tr*_ = {(*x*_*i*_, *y*_*i*_), *i* = 1, …, *N*_*tr*_}, where *y*_*i*_ ∈ *Y*_*tr*_ denote the training kinases, learning process for the zero-shot-learning model involves learning of the compatibility matrix *W* and the BRNN model parameters. Assuming that the training data contain independently and identically distributed samples, we estimate *W* that maximizes the likelihood of observing the training data:

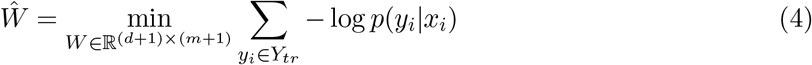

The class posterior probabilities above are provided in Equation (1). Maximizing the likelihood is equivalent to minimizing the cross-entropy loss. We train the model end-to-end by connecting the BRNN model to ZSL model (Figure 3). In this way, the BRNN model learns phosphosite embeddings specifically useful for the ZSL model and classification of kinases. To avoid overfitting, we employ drop-out regularization with a 0.5 keep probability [55]. We apply batch normalization in LSTM cells [56] to normalize the embeddings passed onto the ZSL model. We initialize the *W* matrix randomly from a uniform distribution and minimize the cross-entropy loss function using Adam optimizer [57] with learning rate 10^−4^. The attention weights are also initialized randomly from a normal distribution with a mean of 0 and standard deviation of 0.05. The learning rate and the number of iterations are optimized on validation data (see Section 3.1 for an explanation of the validation data). To reduce the variance of the model, we ensemble 10 models each of which trained with different initializations of the model parameters. The final class probabilities are obtained by averaging output probabilities over the ensemble.

**Figure 3:**
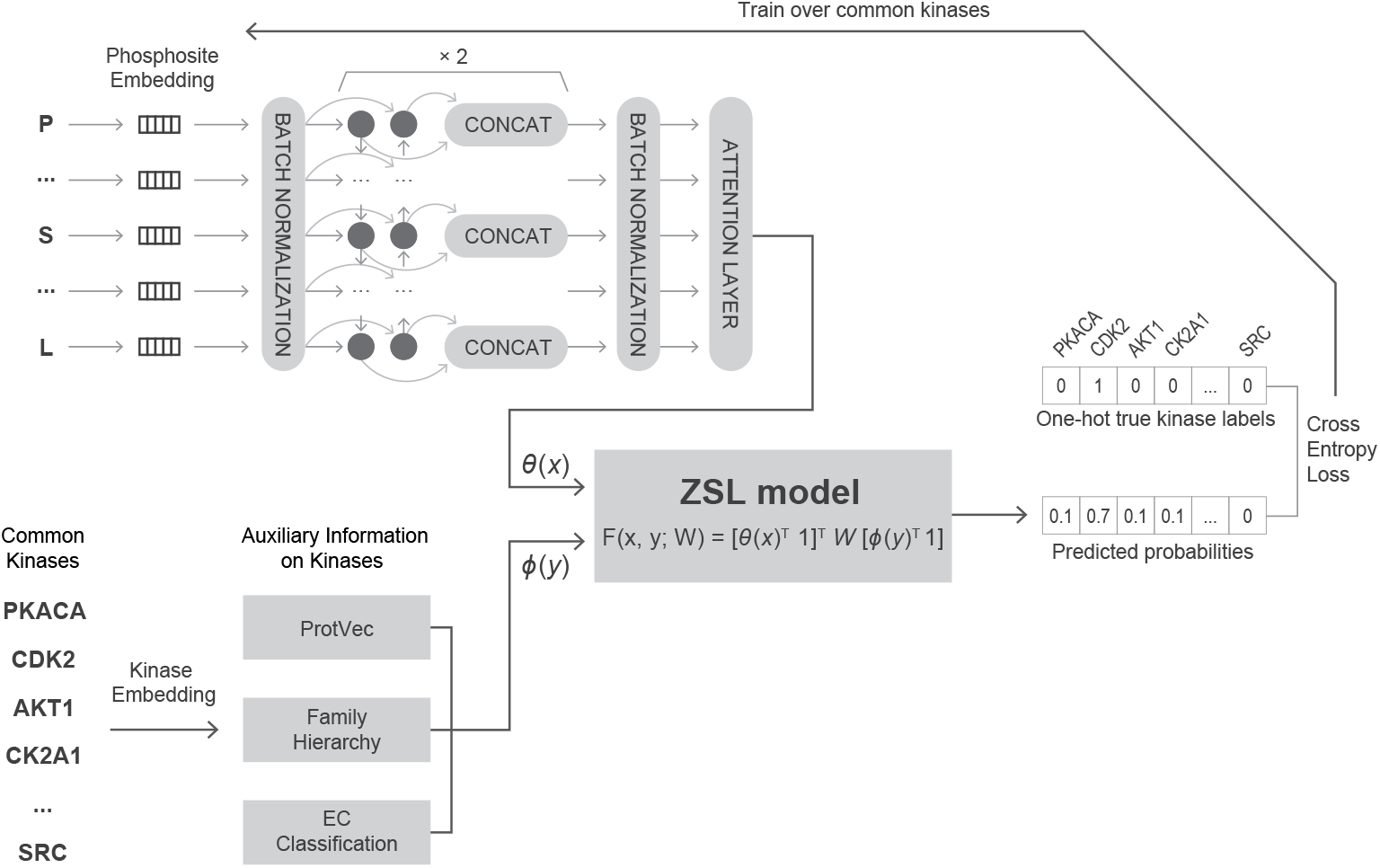
The architecture of DeepKinZero. First, the embedded vectors of phosphosites are passed to a 2-layer bidirectional LSTM network, and then the results after passing through an attention layer are input to the ZSL model. The whole model is trained end-to-end over the common kinases.

#### Making Predictions with the DeepKinZero

The estimated 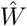 is used at the test time; given a specific input phosphosite, the predicted kinase class, *y*\*, is assigned by maximizing *F* over the test classes:

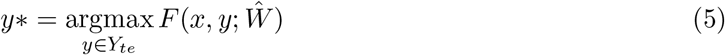

This is equivalent to getting the class with the highest posterior probability as the posterior probability given in Equation (1).

##### 2.2.1 Phosphosite Embeddings

In learning the phosphosite embeddings, we experiment inputting the phosphosite sequence with three different vector representations into the BRNN:

i. **One-hot encoded vector:** Each residue of a peptide sequence is coded with a 21-dimensional vector with binary entries. 20 of these dimensions encode for each of the amino acids and one extra entry is used to encode for non-extant residues. This may happen if the phosphosite is too close to the N-terminal (or the C-terminal) of the protein such that the peptide sequence is shorter than 15 residues. Eventually, with one-hot encoding, each phosphosite sequence is embedded into a 21 × 15 = 315-dimensional binary vector.
ii. **Physical and chemical characteristics of amino acids:** We also use a reduced alphabet that represents each sequence based on the physicochemical properties of the amino acids (AA Prop) in the sequence. We consider the charge, polarity, aromaticity, size, and electronic-property of each amino acid. The categorization of each amino acid into groups based on these five properties are obtained from [58] and is also listed in Supplementary Table 1. Using this categorization, we code each sequence based on property-based one-hot encoded vectors and concatenate them. Charge, size and aromacity properties can each take 3 different values, polarity can take 2 and electronic property can take 5 different values. Therefore, the resulting one-hot encoded vector is 15 × 16 = 240-dimensional.
iii. **ProtVec:** Motivated by the demonstrated success of word embedding techniques in natural language processing (e.g., Word2Vec [59]), unsupervised embedding models have been developed to represent protein sequences, as well. Among these models, ProtVec [60] provides a continuous representation of protein sequences and is trained on sequences from Uniprot-SwissProt using a Skip-gram neural network [61]. ProtVec converts each 3-gram in input sequence into a vector of length 100. There are 13 3-grams in a peptide of 15 residues, thus, our ProtVec representation of each sequence is 13 × 100 = 1300-dimensional.

##### 2.2.2 Kinase Embeddings

The key to zero-shot learning is to know, for each unseen class, the relationship with the formerly seen classes. To establish this relationship between common and rare kinases, we create four different class embedding vectors, which are then concatenated to form a kinase embedding vector, *ϕ*(*y*) in Equation (3). Supplementary Figure 1 summarizes the size of the kinase embedding vectors when all the sources are used. We experiment the utility of some of the vectors through computational experiments and drop those that are not informative in the final model. Below, we give a detailed account of the sources and the way they are deployed to arrive at the desired kinase embeddings:

i. **Kinase hierarchy:** We use the classification proposed by [22]. The data is obtained from Kinase.com [62] (downloaded in June 2018). Supplementary Figure 2 shows this hierarchy. In this classification, there are 10 groups, and 116 families. We convert this to a binary vector by representing families, groups and individual kinases as one-hot encoded vectors. In the end, we attain a binary vector with a size of 583.
ii. **EC classification of kinases:** An alternative source of kinase categorization is the Enzyme Comission(EC) classifications provided by the ENZYME database [63](downloaded in June 2018). According to this classification scheme, kinases are grouped into 6 main categories based on their functions. The two largest categories of kinases are the tyrosine-specific protein kinases and serine/threonine kinases. The main categories are further divided into subcategories (as shown in Supplementary Figure 3).
iii. **Kin2Vec:** As kinases can be related through their kinase domain sequences, we use a ProtVec representation of kinase domain sequences just as we do for the input phosphosite sequence. To differentiate the two, we refer to them as Kin2Vec. ProtVec creates vectors of length 100 for each 3-gram in the sequence and since for each kinase, the kinase domains can be of different lengths, we average the ProtVec vectors generated for each 3-gram into one vector with 100 components.
iv. **KEGG pathways:** To capture the relatedness of kinases in the biological functional space, we create kinase vectors based on the pathways in which the kinases participate. The human pathways are obtained from KEGG database [64–66] (downloaded in April 2018). Cumulatively, there are 190 KEGG pathways in which at least one of the kinases participate. Each kinase vector is formed as a 190-element binary vector based on its participation in each of the cellular pathways.

## 3 Results

### 3.1 Evaluation Protocol

We train and evaluate our models on the experimentally validated kinase-phosphosite associations obtained from the PhosphoSitePlus database [43] (downloaded in March 2018). We exclude iso-form and fusion kinases. The dataset includes 13,426 experimentally identified phosyphorylation sites and their associated 343 kinases. Following the evaluation protocol suggested by Xian et al. [46], we keep the zero-shot kinases well apart from the rest of the classes in learning the models and parameter tuning. We split the data into training, validation and test data based on the number of sites that are associated with each kinase. Kinases with more than 5 sites are considered as training classes. There are 124 such kinases. DeepKinZero is trained on this set, which contains kinase-substrate associations of 12,901 phosphorylation sites with these 124 kinases. The validation set includes the kinase-phosphosite associations of 17 kinases for which there are exactly 5 phosphorylation sites. This validation set includes 80 phosphorylation sites associated with these 17 kinases. The remaining 112 kinases with less than 5 positively labeled examples constitute the test or zero-shot classes. The test data includes these 112 kinases and kinase-phosphosite associations involving 237 phosphorylation sites.

### 3.2 Performance Criteria

To assess the overall performance, we use hit@k accuracy. This metric evaluates performance in terms of the number of times in which the correct class is among the top *k* predicted classes, where *k* is a parameter. If the true class is within the top-k predicted classes, it is considered a true positive prediction. We report results for values of *k*=1,3 and 5. In cases for which a phosphosite is associated with more than one kinase, we consider the prediction to be a true positive if the model predicts one of these kinases for the corresponding phosphosite in the top *k* prediction. In our test dataset, 215 phosphosites are associated with a single kinase, 16 phosphosites are associated with 2 kinases, and 2 phosphosites are associated with 3 kinases. Thus, multi-class instances are rare.

### 3.3 Zero-Shot Learning Results

The representations of the site sequences and the kinases are critical components of the model and they can greatly influence prediction performance. For this reason, we assess the performance of DeepKinZero by comparing the prediction performance of DeepKinZero with different phosphosite and kinase embeddings.

#### 3.3.1 The Effect of Different Phosphosite Representations on Accuracy of Predictions

To thoroughly asses the effectiveness of different phosphosite representations, DeepKinZero is trained with three different input representations: One-Hot, Amino Acid Properties (AA Prop) and ProtVec with and without using BRNN. When a BRNN is employed, the BRNN is trained with the specified site sequence embeddings and the final layer of the BRNN is used as the final sequence embedding and directly input to the zero-shot classifier. Figure 4 summarizes the results of using different phosphosite sequence embeddings. As shown in Figure 4, with respect to hit@1 and hit@3 metrics, the model trained with a BRNN coupled with ProtVec vectors performs the best, where the true kinase is predicted as the top kinase for more than 20% of the sites, and it is among the top 3 for more than 30% of the sites. With respect to hit@5 metric, the input representations have less effect on the prediction performance, where amino acid properties with BRNN delivers the highest hit@5 accuracy with the true kinase being among the top 5 for more than 40% of the sites. Additionally, we observe that the use of BRNN model improves the performance. The model without BRNN embeddings that uses One-Hot sequence embedding as input only returns the true kinase as the top prediction in 10.55% of the test cases. On the other hand, the model with BRNN and ProtVec site embeddings predict the right class with 21.52% accuracy. Note that these numbers are highly impressive since it would not be possible to train predictive models for these kinases due to the inadequacy of training samples, and random guess will achieve only 0.89% accuracy since there are 112 test classes.

**Figure 4:**
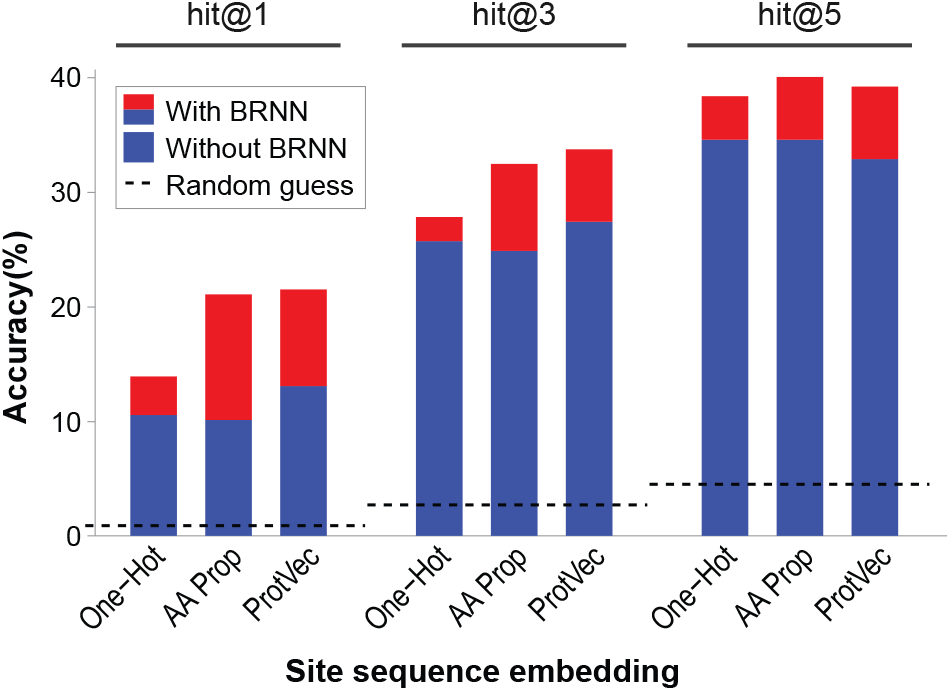
The effect of phosphosite representations on the accuracy of predictions. The hit@1, hit@3, and hit@5 performance of DeepKinZero (percentage of phosphosites for which the top kinase is respectively among the top 1, 3, and 5 predictions) with six different phosphosite embedding methods (one-hot, amino-acid properties, ProtVec, each with or without a Bi-Directional Recurrent Neural Network(BRNN)) are shown. For reference, the hit@1, hit@3, and hit@5 of a random guess (the only existing alternative for the kinases tested) performance are also shown.

To probe the usefulness of the representations learned by BRNN, we use nonlinear dimension reduction. We visualize the BRNN embeddings in a lower nonlinear dimension reduction to visualize the BRNN embeddings in a lower dimensional space using t-distributed stochastic neighbor embedding (t-SNE) [67]. Supplementary Figure 4 shows that the BRNN can separate the examples in the case of kinase groups better than the ProtVec representations, hinting that it successfully captures additional critical information about kinases.

#### 3.3.2 Effect of Kinase Embbeding on Accuracy of Predictions

The performances of models trained with different kinase embeddings is shown in Table 2. In these experiments, for phosphosite embedding, we use a BRNN trained on ProtVec and compare different combinations of class embedding features with each other. To establish a baseline, the first row shows the accuracies attained using a random guess. The second row lists the performance of the model when we input the one-hot vector of kinases as class embeddings; this model is effectively a model that does not transfer knowledge between different kinases. As shown in the table, the performance of this model is worse than a random guess, demonstrating that learning is non-trivial if the class embeddings are not included.

**Table 2:** Effect of kinase embedding on the accuracy of predictions. The hit@1, hit@3, hit@5, and hit@10 performance of DeepKinZero using all possible combinations of four different kinase embeddings are shown. Each row shows a model with a specific combination of kinase embeddings, where the check marks indicate that the corresponding kinase embedding is included in the model. For reference, the performance of random guess and an embedding that only uses the identity of individual kinases (thus does not transfer information between kinases) is also shown.

| Family Hierarchy | Pathways | EC Classification | Kin2Vec | hit@1 | hit@3 | hit@5 | hit@10 |
| --- | --- | --- | --- | --- | --- | --- | --- |
| Random Guess |  |  |  | 0.89 | 2.70 | 4.50 | 9.30 |
| One-hot vector of kinases as class embedding |  |  |  | 0.84 | 1.69 | 2.95 | 7.59 |
| ✓ |  |  |  | 17.72 | 31.65 | 37.55 | 46.84 |
|  | ✓ |  |  | 8.02 | 13.5 | 16.03 | 21.52 |
|  |  | ✓ |  | 5.06 | 13.5 | 17.72 | 30.8 |
|  |  |  | ✓ | 1.27 | 5.91 | 8.02 | 16.03 |
| ✓ | ✓ |  |  | 14.77 | 27.85 | 35.02 | 46.84 |
| ✓ |  | ✓ |  | 19.41 | 29.96 | 38.82 | 47.68 |
| ✓ |  |  | ✓ | 18.99 | 33.33 | **40.08** | 47.26 |
|  | ✓ | ✓ |  | 8.86 | 11.39 | 19.41 | 28.69 |
|  | ✓ |  | ✓ | 8.02 | 13.5 | 16.46 | 21.94 |
|  |  | ✓ | ✓ | 6.75 | 14.77 | 19.83 | 32.49 |
| ✓ | ✓ | ✓ |  | 15.19 | 25.74 | 35.86 | 46.84 |
| ✓ | ✓ |  | ✓ | 15.61 | 30.38 | 36.29 | 45.99 |
| ✓ |  | ✓ | ✓ | **21.52** | **33.76** | 39.24 | 47.68 |
|  | ✓ | ✓ | ✓ | 10.55 | 18.14 | 24.05 | 32.91 |
| ✓ | ✓ | ✓ | ✓ | 16.88 | 29.11 | 34.18 | **48.10** |

The next four rows in the table show the results of the models trained with kinase embedding vectors of individual data sources. Thus, they portray the strength of each source in isolation from the others. Among the four possible kinase embeddings, the kinase hierarchy is the leading contributor to the accuracy of the model, achieving 17.72% accuracy when used as the sole auxiliary information source. As this hierarchy reflects the functional and evolutionary information (based on sequence similarities) on the kinases, it is expected that they carry valuable information about kinase similarities. When used in isolation of other sources, Kin2Vec is found to be the least useful source.

The next set of results display the combinations of two sources. In all classes, combining family hierarchy with another information improves the model’s performance the most. The model achieves 18.99% hit@1 accuracy by combining family hierarchy with Kin2Vec. Furthermore, combining family hierarchy with EC classification or Kin2Vec vectors increases hit@5 accuracy from 37.55% to 38.82% and 40.08% respectively. Also among all combinations, its removal from the model affects the accuracy most adversely(for example second to the last row in the table).

Overall, the best performance is achieved by using family hierarchy, EC classification and Kinase2Vec vectors which achieves 21.52% on hit@1 accuracy, 33.78% on hit@3 and 39.24% on hit@5 accuracy. Adding pathway vectors into this combination degrades the hit@1, hit@3 and hit@5 accuracies significantly, although the use of pathways alone is the second best (fourth row) when used individually as an embedding and it improves the hit@10 accuracy. It is possible that the information provided by pathway membership may not be sufficiently specific to contribute additional information on the relationships between kinases. When hit@5 or hit@10 is used, all the models except those that ignore the family hierarchy performs relatively well. The best performance is achieved when all the available information is included in the model (48.1 %).

#### 3.3.3 Inspecting Model Weights

We further analyze the learned weights in the model to gain further insight into the model. First, we inspect BRNN attention weights. Figure 5 a) shows the average attention assigned to each position in the input sequence by the BRNN model. The center residue emerges as the most important residue, thus the model correctly learns to assign more weight to the center, where the phosphosite is located at. The immediate neighbors and the residues within 2 positions are the next most important residues. This aligns well with our expectations.

**Figure 5:**
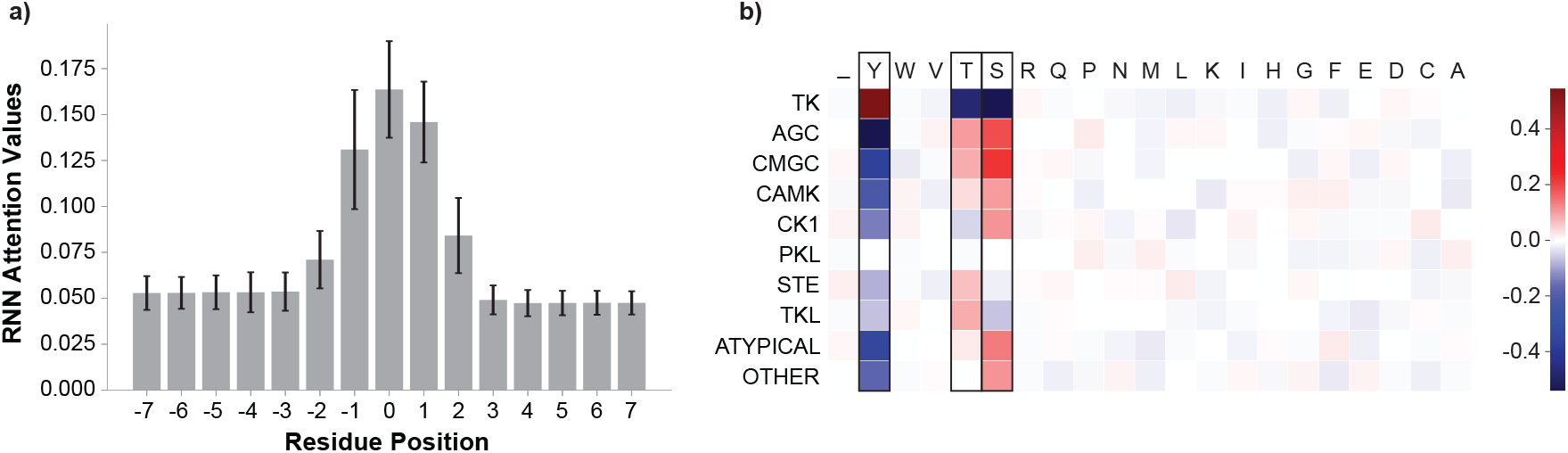
Position and amino acid type weights. (Best viewed in color) a) Average attention weights of the residue positions calculated over the ensemble BRNN model. Residue position 0 is the phosphosite position. b) Average zero-shot learning weights for each amino acid type at the phosphosite.

Next, we investigate the amino acid type importance at the phosphosite. Recall that the *W* matrix specifies the relative contribution of the correspondence between each dimension in the kinase embedding space with each dimension in the site embedding space. To investigate the weights assigned to each amino acid type at the phosphosite embedding, we calculate the average weights assigned to different amino-acid types for each group of kinases at the phosphosite. As clearly seen in Figure 5 b) S, Y and T correctly receive the largest weights. Moreover, the weights assigned to a different type of amino acids in each group align well with existing knowledge of kinase groups. For example, the TK family, which exclusively works on thyrosine residue (Y), puts a very large positive weight on tyrosine while other families do not. Similarly, CMGC work predominantly on serine (S) and threonine(T) and these are the two residues that get a positive large weight. PKL group is a diverse group that could be the reason why neither of the residue types emerges as predominantly predictive.

#### 3.3.4 Validation on an External Data

We also evaluated DeepKinZero on an external test data we had retrieved from PhosphoELM database [68] (downloaded on September 2018). We first removed all the kinases and their associated phosphosites that were in our training and validation set from PhosphositePlus dataset. The remaining kinase-substrate associations in the PhosphoELM dataset represent an instance that is well-suited to DeepKinZero’s objective, in that almost all of the kinases in this dataset have very few known associations. To be more precise, there are 52 phosphosites associated with 40 kinases and 29 of these kinases have only one site associated with it. One of them have 7 sites associated with it, while the other 10 kinases have 4, 3 or 2 associated sites. DeepKinZero trained on PhopsphositePlus and evaluated on this PhosphoELM dataset achieves hit@1 accuracy of 33.96%, hit@3 accuracy of 52.83%, 62.26% hit@5 accuracy and 77.36% hit@10 accuracy. Although the dataset is small, it provides confidence that the model generalizes to other datasets.

#### 3.3.5 Comparison with Other Methods

In the literature, there are no models that we can directly compare our method against. However, there are two methods [69, 70] that aim at a different but a related problem. These two methods are designed to predict the phosphosites for kinases with no known sites, which is the reverse scenario of our problem; we predict the kinase of a given phosphosite. Predikin [69] operates with a set of rules governing the amino acids around the phosphosites. These rules, however, are derived from 3D structures of kinases bound to their substrates. Therefore, the method is limited by the availability of the protein structures and cannot be applied to kinase families without 3D structures. Because the Predikin server was not available, we were not able to carry out a comparison with this method.

The second method is by Wagih et al. [70], which is based on the idea that, as compared to a random set of proteins, interaction partners of a kinase are more likely to be phosphorylated by that kinase. Thus, the method finds enriched motifs in the interaction partner sequences to predict sequences that a kinase can bind. The method is not applicable, however, when the kinase has a low number of interaction partners and/or the number of phosphosites on the interactors is low. Our method predicts the kinase of a given phosphosite, whereas Wagih et al. predicts the phosphosite of a kinase. Thus, the two methods are not directly comparable but still, we conduct the following comparison. For the 112 zero-shot kinases, we predict the motifs by Wagih et al. model. If we consider the top motif returned, the method correctly matches 11 of the phosphosites of the 112 kinases, leading to 9.8% hit@1 accuracy. If we consider the top 5 motifs returned for each kinase, the correct phosphosite sequence matches 26 of phosphosites of the 112 kinase motifs leading to 23% hit@5 accuracy. These numbers are significantly lower than what DeepKinZero can achieve (21.52% and 40.08%). We should note that this comparison also favors Waigh et al. because DeepKinZero is evaluated based on how many phosphosites it gets right from all the available phosphosites. This is twice the number of kinases over which Waigh et al. was evaluated with.

## 4 Conclusion and Future Work

Many kinases are understudied with no known target proteins or sites; therefore, only a small subset of kinases dominates the annotated phosphosite databases. DeepKinZero, unlike conventional supervised methods can offer predictions for kinases which do not have any known phosphosites. The zero-shot learning framework transfers knowledge from common kinases to rare kinases and this way it renders the predictions for classes that were never observed in the training phase possible. Exploring the lesser-studied kinases and their associated substrates and sites will likely reveal major insights into the healthy and diseased cell. Through guiding experimental studies, we hope DeepKinZero will help in illuminating the dark phosphoproteome.

The work presented here can also be extended in different dimensions, which we plan to study in our future work. First of all, the ability to transfer learning between classes is based on the ability to define the characteristics of the kinases as vectors, which is derived from auxiliary information on kinases, such as taxonomies of kinases or deep representation of their kinase domain sequences (as detailed in Section 2.2.2 section). These kinase embeddings can be extended to include further information. For a kinase to catalyze a phosphorylation event on a substrate, peptide specificity on the substrate is considered as the main determining factor. However, the peptide specificity is not the only element. The cellular localization and the structural domains outside the catalytic domain have also been reported to be important factors. Thus, in deriving the kinase embeddings, other information such as cellular localization can be used.

A second line of work is to extend this work to general zero-shot learning. The zero-shot learning assumes that the testing instances are only classified into the candidate unseen classes. In this study, we also assume that the candidate classes at the time of testing all belong to the rare kinases. The generalized zero-shot learning is a more open setting where all the classes (seen and unseen) are available as candidates for the classifier at the testing phase [71]. This is a much harder problem due to the greater number of classes handled during testing. Additionally, the classifier tends to assign instances into one of the previously exposed classes. This problem needs more specific methods. In future work, we plan to extend this framework to handle this generalized setting.

## 5 Acknowledgements

This work was supported by internal fundings of Sabanci and I.D. Bilkent Universities.

## 6 Author contributions

I.D., M.K. and O.T. designed the study and wrote the manuscript. I.D. implemented the model, I.D. and B.A. curated datasets and conducted the computational experiments.

