## Supplementary Tables and Figures for "DeepKinZero: Zero-Shot Learning for Predicting Kinase-Phosphosite Associations Involving Understudied Kinases"

| AA | Charge | Polarity | Aromaticity | Size | Electronic Property |
|---|---|---|---|---|---|
| A | Neutral | Non-polar | Neutral | Small | Strong Donor |
| R | Positive | Polar | Neutral | Large | Strong Acceptor |
| N | Neutral | Polar | Neutral | Medium | Strong Acceptor |
| D | Negative | Polar | Neutral | Medium | Strong Donor |
| C | Neutral | Polar | Neutral | Large | Neutral |
| Q | Neutral | Polar | Neutral | Large | Weak Acceptor |
| E | Negative | Polar | Neutral | Large | Strong Donor |
| G | Neutral | Non-polar | Neutral | Small | Neutral |
| H | Positive | Polar | Aromatic | Large | Neutral |
| I | Neutral | Non-polar | Aliphatic | Large | Weak Donor |
| L | Neutral | Non-polar | Aliphatic | Large | Weak Donor |
| K | Positive | Polar | Neutral | Large | Strong Acceptor |
| M | Neutral | Non-polar | Neutral | Large | Weak Acceptor |
| F | Neutral | Non-polar | Aromatic | Large | Weak Acceptor |
| P | Neutral | Non-polar | Neutral | Small | Strong Donor |
| S | Neutral | Polar | Neutral | Small | Neutral |
| T | Neutral | Polar | Neutral | Medium | Weak Acceptor |
| W | Neutral | Non-polar | Aromatic | Large | Neutral |
| Y | Neutral | Polar | Aromatic | Large | Weak Acceptor |
| V | Neutral | Non-polar | Aliphatic | Large | Weak Donor |

Table 1: Classification of amino acids(AA) based on five different physiochemical amino acid properties as in [1].

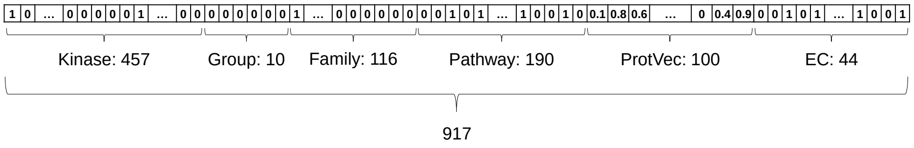


Figure 1: The one-hot encoding representation of the class embeddings. Vectors from different sources are concatenated to form the class embedding vector. The numbers state the size of the vectors culled from each data source. The number 917 is the total size of the class embedding vector.

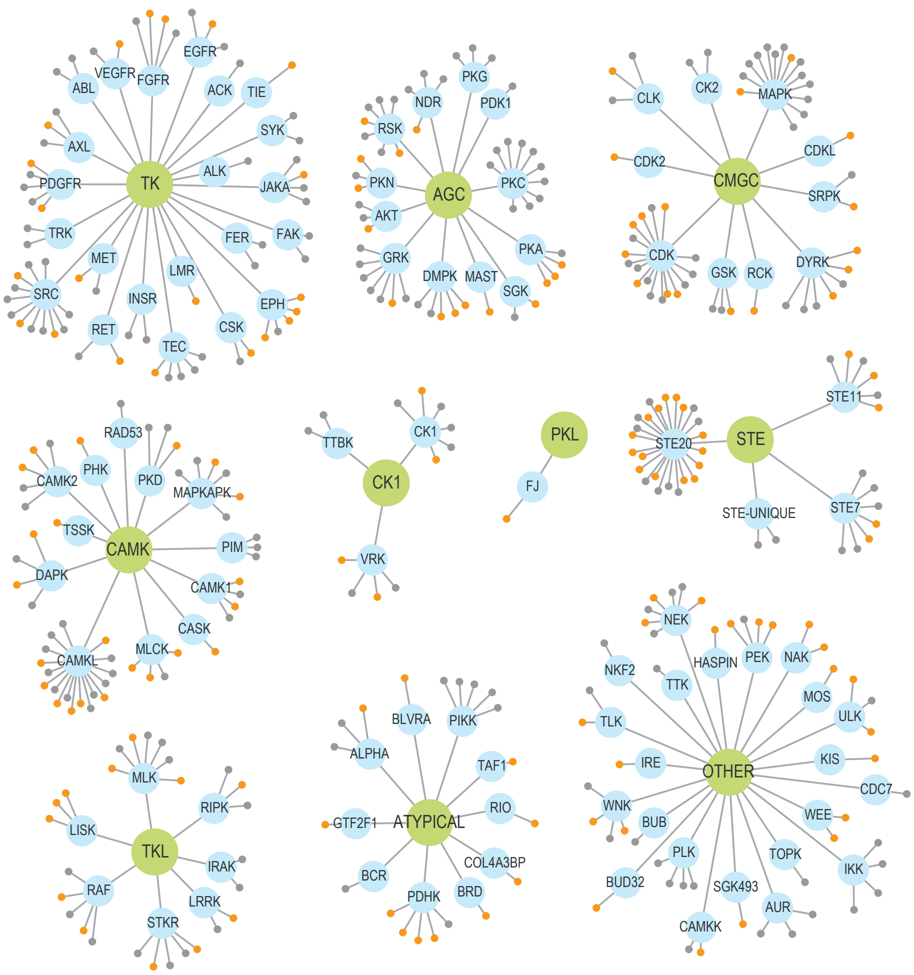


Figure 2: The partitioning of kinases into families and groups as proposed in [2]. Each network is centered on a group node shown in green. The families (blue nodes) that are listed under a group are linked with edges to the group node. The small gray nodes show kinases with many known sites; these kinases are used in the training of the models while the small orange nodes indicate the kinases that are unseen and used for testing.

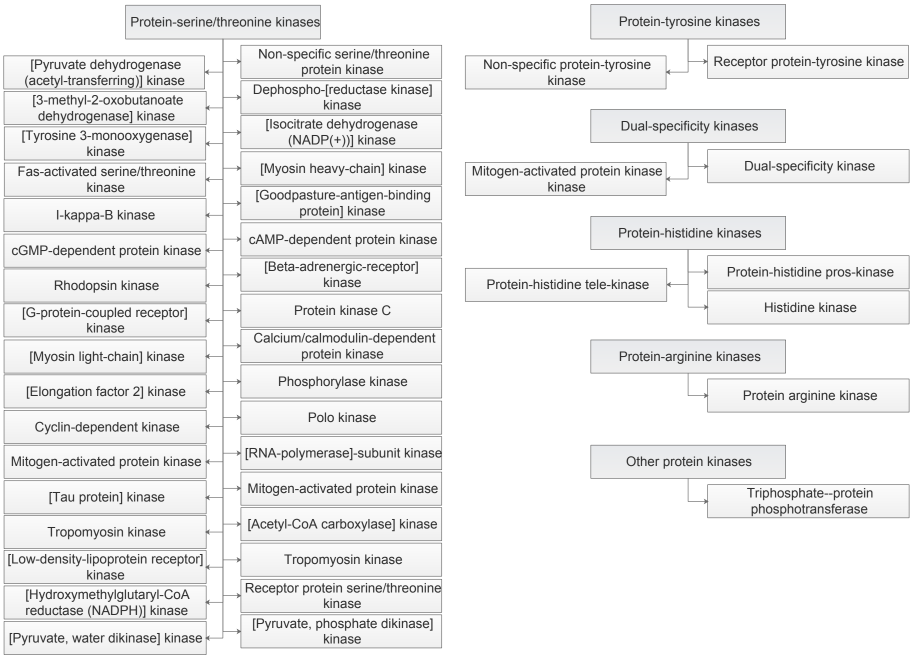


Figure 3: Classification of kinases according to the ENZYME database.

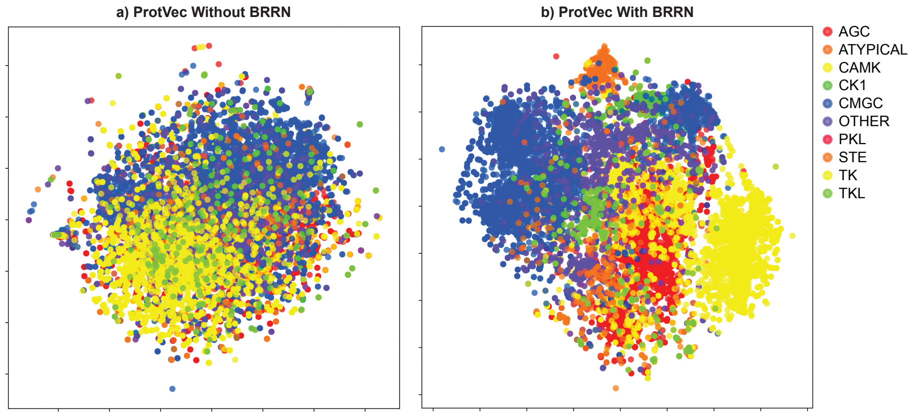


Figure 4: t-SNE representation of the BRNN embeddings generated with and b) without BRNN on ProtVec phosphorylation site representation. The colors represent different kinase groups.
